# Fine-scale flight behaviour reveals eagles’ response to different uplift sources and highlights observational gaps in high-resolution weather models

**DOI:** 10.64898/2026.08.18.745477

**Authors:** Francesca Frisoni, Tom Carrard, Martin U. Grüebler, Julia S. Hatzl, Kamran Safi, Michael A. Sprenger, Petra Sumasgutner, Martin Wikelski, Martina Scacco

**Affiliations:** Department of Migration, Max Planck Institute of Animal Behavior, Radolfzell, Germany; Department of Biology, University of Konstanz, Konstanz, Germany; Institute for Atmospheric and Climate Science, ETH Zürich, Zürich, Switzerland; Swiss Ornithological Institute, Sempach, LU, Switzerland; Department of Landscape Ecology, ETH Zürich, Zürich, Switzerland; Konrad Lorenz Research Center (KLF), Core Facility for Behavior and Cognition, Grünau/Almtal, Austria; Department of Behavioural and Cognitive Biology, University of Vienna, Vienna, Austria; Dynamic Macroecology, Swiss Federal Institute for Forest, Snow and Landscape Research WSL, Birmensdorf, Switzerland

**Keywords:** discriminant function analysis, uplift classification, opportunistic behaviour, vertical wind velocity, animal borne sensors, machine learning, gravity waves

## Abstract

Understanding how animals respond to their physical environment requires environmental observations at the scale at which behavioural decisions are made. For soaring birds, the coarse resolution of weather products has long hindered the analysis of their behavioural response to fine-scale atmospheric dynamics, forcing uplift sources to be inferred largely from behaviour itself. Here, we combined high-resolution movement data from 24 golden eagles with the kilometre-scale COSMO weather model. We first classified thermal, orographic, and gravity-wave uplifts using independent atmospheric predictors and then quantified the birds’ use of each uplift type and their fine-scale behavioural responses. Eagles relied predominantly on thermals, but opportunistically adjusted their use of uplift sources seasonally. The birds’ flight behaviour could not reliably indicate which uplift type was primarily used, and thus suggests that both atmospheric processes and behavioural responses are better described as continua than discrete categories. Finally, we compared vertical wind velocities derived from eagles soaring behaviour with those modelled by the COSMO weather model, showing that most of the thermals exploited by eagles remain unresolved at kilometre-scale model resolution. Our results demonstrate how high-resolution weather models provide new insights into bird movement decisions, while also highlighting the potential of soaring birds as biologically embedded atmospheric sensors that could help closing the resolution gap in atmospheric models.

## 2 Introduction

The movement cost of flying animals is dependent on the fine-scale dynamics of the atmosphere, which indirectly impact movement decisions, large-scale movement patterns and fitness (Kunz et al. 2008, Shepard, Ross & Portugal 2016, Hurme et al. 2025). Soaring birds are the epitome of this dependency: their movement costs and decisions are determined by the availability, distribution, and quality of rising air currents (uplifts). As uplift support can reduce flight costs by nearly 80%, bringing them close to resting metabolic levels (Duriez et al. 2014, Harel et al. 2016, Williams et al. 2020), soaring birds represent an ideal system to investigate fine-scale behavioural responses to such dynamic physical environments.

Soaring flight is traditionally understood to be sustained by two main categories of uplifts: convective (thermals) and dynamic uplifts (mechanically driven uplifts such as orographic uplift and mountain gravity waves) (Kerlinger 1989, Garstang et al. 2022, Carrard et al. 2025). Convective and dynamic uplifts differ in their formation, spatio-temporal distribution and physical properties, such as stability and strength (Organisation Scientifique et Technique Internationale du Vol à Voile 2009). Thermals are ephemeral ”bubbles” of rising warm air, produced by uneven heating of the terrain surface (Akos et al. 2010). Dynamic uplifts are mechanically induced by the deflection of horizontal winds off the underlying topography such as mountain ridges (Brandes & Ombalski 2004, Etling 2014, Bohrer et al. 2012). Such wind deflections produce an ascending orographic uplift on the windward side of a slope, and, depending on the wind speed and the stratification of the atmosphere, can lead to the formation of horizontally propagating mountain gravity waves on the leeward side (Durran 1990, Whiteman 2000). While thermals are usually characterised by high vertical wind velocities, they have limited spatial extent, are patchily distributed, can be highly turbulent at their margins and are tilted and weakened by wind shear (Organisation Scientifique et Technique Internationale du Vol à Voile 2009). In contrast, dynamic uplifts generally cover larger spatial extents and are characterised by more predictable but lower vertical wind velocities (Brandes & Ombalski 2004, Etling 2014). The different physical properties of uplifts, coupled with observations of soaring birds’ behaviour, led to a long-standing hypothesis in the field: namely that soaring birds would adopt distinct flight modes depending on the uplift source they were using. It was assumed that circular, spiralling flight would reflect the use of thermals, and linear soaring flight the use of dynamic uplifts (Alerstam & Hedenström 1998, Kerlinger 1989, Santos et al. 2017). However, while this hypothesis found support under specific conditions (Santos et al. 2017), its general validity has remained largely and quantitatively untested, due to the lack of atmospheric data at the resolution required to actually evaluate it (Santos et al. 2017, Treep et al. 2016, Shepard, Williamson & Windsor 2016, Shamoun-Baranes et al. 2016, Scacco et al. 2019).

Characterising individual uplifts requires atmospheric data at scales of minutes and hundreds of metres. Therefore, publicly available weather products such as ERA5, providing hourly data at 31 km resolution (Hersbach et al. 2023), cannot resolve single uplifts nor distinguish between their sources (Murgatroyd et al. 2018, Carrard et al. 2025). In contrast, with the advancements in biologging technology, quantitative high (sub-second) resolution behavioural data are now available widely. While this mismatch in resolution between meteorological and behavioural data has long hindered our ability to investigate the behavioural plasticity of soaring birds under different atmospheric conditions (Murgatroyd et al. 2018, Hanssen et al. 2020), it also points to an underexplored opportunity: behavioural data encode atmospheric information at scales weather models cannot resolve, and soaring birds may themselves constitute a distributed observation network for fine-scale atmospheric dynamics. The extraction of atmospheric parameters such as uplift strength and horizontal wind speed from birds’ flight trajectories (Treep et al. 2016, Weinzierl et al. 2016, Goto et al. 2017) has already been explored in the literature, but once again validation remained mostly hindered by the resolution of the meteorological data used. Resting models and validations on coarse synoptic conditions and observational proxies therefore created a circularity in the analysis of behavioural responses: atmospheric dynamics conditions inferred from observed behaviour, and behaviour interpreted through assumed atmospheric dynamics (Kerlinger 1989, Treep et al. 2016).

Recent advancements in meteorological modelling are finally enabling atmospheric representations at scales of 1 to few kilometres (Benjamin et al. 2016, Zängl et al. 2015, COSMO Consortium 2024, Prein et al. 2026) and have begun to open this gate. While still insufficient to fully capture the atmospheric context experienced by birds at the fine-scale at which their flight behaviour is recorded, these models hold promise for a deeper understanding of the spectrum of soaring birds’ behavioural responses to fine-scale atmospheric dynamics and of the potential of these birds as moving sensors of atmospheric conditions. A first study exploiting this resolution quantitatively and systematically demonstrated that golden eagles regularly use gravity waves in the Alps, and already challenged the simple association between flight mode and uplift source (Carrard et al. 2025); the generality and quantitative scope of this challenge, however, remain to be established.

Here, we combine sub-second behavioural movement data from 24 golden eagles (*Aquila chrysaetos*), collected with GPS and inertial measurements units (IMU), with the high-resolution regional weather forecasting model COSMO (COSMO Consortium 2024), to investigate the behavioural response of soaring birds to atmospheric dynamics at an unprecedented resolution. The spatial extent of the tracking dataset and the COSMO model cover the European Alps and Apennines, regions of complex topography where uplifts of different sources are abundant and co-occur. This rich dataset allows us to ask three central questions: (i) Do eagles use the different available uplift sources in proportion to their environmental availability, and how does this pattern vary across seasons? (ii) Do they adjust their flight behaviour depending on different uplift sources such that behavioural data alone could reliably inform us about the uplift source powering a soaring event? (iii) Beyond categorical uplift classification, what do eagles’ climbing rates reveal about the fine-scale vertical wind velocities that kilometre-scale weather models cannot resolve? Using an existing training dataset of 209 manually labelled uplift sources (Carrard et al. 2025) we: 1. Built an automatic classification algorithm for the identification of uplift sources based only on weather and topographic parameters; 2. Applied this algorithm to characterise and compare the uplift used by the 24 eagles, with those available in the environment, at the times when eagles were flying; 3. Calculated 59 behavioural metrics from GPS and IMU and use a machine learning algorithm to test whether uplift sources can be differentiated from a behavioural perspective; 4. Compared uplift strength (vertical wind velocities) derived from eagles’ soaring behaviour with those predicted by COSMO, to identify observational gaps in kilometre-scale weather model resolution and assess the potential of movement-derived measurements to help close them.

## 3 Materials and Methods

### Biologging data

#### Data availability

We used Global Positioning System tracking data of 24 individuals, which is part of a large long-term study on juvenile golden eagles in the European Alps and Apennines. The complete dataset is deposited on Movebank.org (Wikelski et al. 2024) (Movebank study name ”Life Track Golden Eagles Alps”) and includes data collected with high-resolution solar GSM-GPS-IMU loggers (e-obs GmbH, Munich, Germany) from 2017 to date. For the purpose of this study we only used a subset of the data corresponding to the years 2020 and 2023, for which COSMO weather data were available. The loggers were fitted on nestlings using a leg-loop harness (weight of harness and logger maximum 60 g, see Zimmermann et al. (2026) for details on fieldwork procedure). The loggers have a regular sampling frequency of one GPS location every 20 min, but with fully charged battery, they collect high-resolution ’super bursts’ at 1 Hz (one location per second) for 5 minutes, every 15 minutes. Concurrently with the super bursts, the loggers collect IMU data (inertial measurements units including accelerometer, gyroscopes, and magnetometer) in bursts of 8 seconds at 20 Hz. GPS and IMU data were collected with an average time lag of 0.3 seconds. For the analysis, we only retained super bursts for which both GPS and IMU were recorded simultaneously at high resolution, resulting in 1,695,418 GPS observations recording over 471 hours of flight.

### Characterisation of soaring behaviour

We defined as soaring (climbing) segments flight bouts in which eagles were gaining height. These were classified as consecutive GPS observations with a positive climbing rate (difference in altitude between consecutive locations, divided by their time lag), smoothed over a moving window of 21 seconds (Carrard et al. 2025). We retained soaring segments with a minimum duration of 7 seconds and assumed shorter segments to reflect small-scale turbulence (in larger species, the duration of one climbing circle was estimated to be 15 seconds (Weinzierl et al. 2016)). The final dataset consisted of 25,173 soaring segments, representing the unit of the behavioural classification analysis. Each soaring segment was associated to the IMU data closest in time, with a time tolerance of 30 seconds.

We characterised soaring behaviour at fine-scale using a total of 59 behavioural parameters, extracted from both GPS and IMU data. Parameters were extracted for each observation (or in some cases calculated between consecutive observations), and then summarised per segment in terms of average, cumulative sum, minimum, maximum, variance and standard deviation. From GPS data we extracted and calculated: flight height of the eagle (above ellipsoid and ground), horizontal displacement between consecutive locations, ground speed (horizontal displacement divided by time lag), vertical displacement, vertical speed (or climbing rate: vertical displacement divided by time lag), cumulative turning angle (cumulative sum of the angles between consecutive locations along the soaring segment), number of changes in headings, and tilt. For ”circular segments” (segments with a cumulative turning angle *≥* 360°) we additionally calculated the number of circles and circling rate, that is the number of circles performed per unit time. From the accelerometer, we extracted the overall and vectorial dynamic body acceleration (ODBA and VeDBA) as proxies of energy expenditure (Wilson et al. 2020), and the standard deviation of the acceleration over the z-axis (vertical ACC axis), as a proxy of flight turbulence (Laurent et al. 2021). From the combination of accelerometer and gyroscope, we extracted the Euler angles of yaw, pitch and roll, describing fine changes in the body posture of the eagle (Safi et al. 2025). As above, we summarised these variables by segment ID. See Supplementary Material SM section S1 for detailed definitions of the behavioural parameters.

### Selection of background points to characterize uplift availability

In order to investigate eagles’ potential selection of specific uplift sources, we compared their use with their availability in the environment. Because individual eagles fly in different regions relative to each other, availability was assessed for each individual at the spatio-temporal scale of its movement. Specifically, for each individual we extracted a convex hull (minimum convex polygon) around all soaring location of that individual across the entire year. Within each individual’s convex hull, we (i) randomly sampled a number of background points five times larger than the number of soaring locations for that individual, and (ii) repeated the same locations for all the days in which the respective individual was flying at their median flight hour. This ensured a good representation of the full range of atmospheric conditions accessible to the individuals at the time when they were flying. This procedure produced a dataset of 144,800 background points between January and December 2023. Each soaring location and each background point were then annotated with topographic and weather parameters.

### Environmental data

#### Topographic parameters

We extracted topographic variables from a digital elevation model (DEM) openly available for Europe at 30 m spatial resolution (Hengl et al. 2020) and used it to extract slope (steepness angle), aspect (orientation of the slope relative to the North) and roughness (a measure of heterogeneity in elevation) layers (Hijmans 2023).

### Weather parameters

Weather parameters were extracted from the regional forecasting model COSMO-1 (Consortium for Small-Scale Modelling, COSMO Consortium (2024)), provided and operationally run by the Swiss national weather service MeteoSwiss (MeteoSwiss 2024) and available to us for the years 2020 and 2023. Specifically, we extracted the following raw variables: horizontal (U and V) and vertical (W or vertical wind speed) wind vector components, air pressure, air temperature, and surface sensible heat flux (a measure of heat flow between the Earth’s surface and the atmosphere, due to temperature differences). From these, we derived additional parameters considered useful proxies to distinguish the atmospheric processes that characterise the different uplift types: horizontal wind speed, orographic lifting coefficient, and value and height of maximum stability using the Brunt–Väisälä frequency (see SM section S2 for details on the calculation).

The vertical wind speed (W) is a direct indication of an uplift, defined as a positive (rising) vertical air flow. The surface sensible heat flux is used as indirect proxy of thermals occurrence, as it becomes large in conditions with strong radiative surface heating, and helps identifying the potential convective source of an uplift even when the short duration or small spatial extent of individual thermals prevents them from being explicitly resolved by the model. The orographic lifting coefficient is a proxy used to indirectly estimate the availability of orographic uplifts (Bohrer et al. 2012), computed using horizontal wind speed and an updraft coefficient that depends on slope angle, terrain aspect and wind direction. Finally, the Brunt–Väisälä frequency is a measure of the static stability of the boundary layer and lower troposphere: higher values indicate a more strongly stratified atmosphere, in which vertical buoyant motions are suppressed while conditions become more favourable for the generation and propagation of mountain gravity waves, provided that sufficient horizontal winds are present; therefore high values are used as indirect indicator that mountain gravity waves are more likely to occur whereas thermal convection is less likely (Durran 1990, Wurtele et al. 1996, Stull 2012).

All soaring and background observations were annotated with the above environmental variables using trilinear interpolation in space and linear interpolation in time. Soaring points below the lowest model level (about 10 m above ground) were annotated with the values of the weather variables corresponding to the lowest model level. For 12 segments, the wind fields could not be annotated due to missing values in the DEM and were therefore excluded. This resulted in a subset of 25,161 soaring segments used in the following analyses. Soaring locations belonging to the same soaring segment (consecutive soaring locations 1 second apart) were considered part of the same uplift event. Therefore as for the flight metrics, also the environmental variables were averaged per segment ID soaring. To decrease computation time, we annotated every second location per soaring segment (every third for segments longer than 10 seconds).

### Automated classification of uplift sources based on environmental parameters

#### Training dataset

Our classification algorithm relied on an existing dataset of 150 labelled uplift sources, produced by Carrard et al. (2025). Because soaring behaviour is only possible in the presence of uplifts, the occurrence of this behaviour can be used to infer uplift presence (Scacco et al. 2019). The training dataset created by Carrard et al. (2025) applied this concept on the same eagles’ dataset used in this study: the authors first randomly extracted 150 eagles’ soaring segments, indicative of uplift presence, and then manually labelled them with the most likely source powering them. The manual labelling was done by expert meteorologists, based on inspection of each soaring segment visualised within their atmospheric context using a series of vertical cross sections. The atmospheric context was quantified using several predictors that capture conditions favourable to each specific uplift source at the synoptic and mesoscale level (see below), extracted from COSMO (COSMO Consortium 2024). The resulting labelled dataset included 72 clearly classified uplifts (7 orographic uplifts, 57 thermal uplifts, and 8 gravity waves), 45 uplifts originating from mixed sources (30 thermal/orographic, 15 thermal/wave) and 33 unknown cases (where weather parameters at the scale offered by COSMO were not indicative of any of the three sources, i.e. low orographic uplift, low surface sensible heat flux and absence of wave). In order to balance the proportion of different uplift sources in our training dataset, we manually labelled additional 225 soaring events, following the same procedure as described in Carrard et al. (2025). This dataset consisted of 63 orographic uplifts, 59 thermal uplifts, 60 gravity waves, 34 mixed sources (19 thermal/orographic, 15 orographic/wave) and 9 unknown cases. From these two datasets we retained only clearly classified uplift sources (excluding unknown cases and mixed events) and excluded segments shorter than 7 seconds. The final dataset consisted of a total of 209 clearly classified uplifts - 57 orographic uplifts, 97 thermal uplifts and 55 gravity waves - that we used as training dataset in the following analysis.

### Discriminant Function Analysis

In order to automatically classify the uplift sources used by the eagles, we used the manually labelled dataset to train a DFA machine learning classification algorithm (using Venables & Ripley (2002)). To evaluate model performance, we applied a stratified 10-fold cross-validation (Kuhn & Max 2008). Specifically, the labelled dataset (n=209 segments) was partitioned into 10 folds, stratified by uplift type (orographic uplifts, thermal uplifts and gravity waves) to preserve the relative proportion of classes across folds. In each of the ten iterations, one fold (10% of the data, on average 20 segments) was held out as test dataset, while the other nine folds (90% of the data) were used as training dataset. This procedure ensured that each segment was included in the test dataset only once across the ten iterations, avoiding uneven sampling and offering complete coverage of the dataset. We used as predictors the atmospheric and topographic proxies described above, averaged along each segment. Namely: surface sensible heat flux, horizontal and absolute value of vertical wind speed, value and height of maximum stability, orographic lifting coefficient, height above ground, slope angle, aspect and roughness. We pooled predictions across all ten iterations into an aggregated confusion matrix (Figure 1). The models were evaluated in terms of overall accuracy (ratio between correctly classified and total uplifts) and intra-class accuracy (ratio between correctly classified and total uplifts for each uplift class) using the model confusion matrix. The DFA models had an overall accuracy of 0.88, with 47/57 orographic uplifts, 96/97 thermal uplifts, and 41/55 gravity waves correctly classified (see SM section S3 and Figures S1-S2-S3 for visual comparison of environmental proxies distributions across manually and automatically classified datasets).

**Figure 1:**
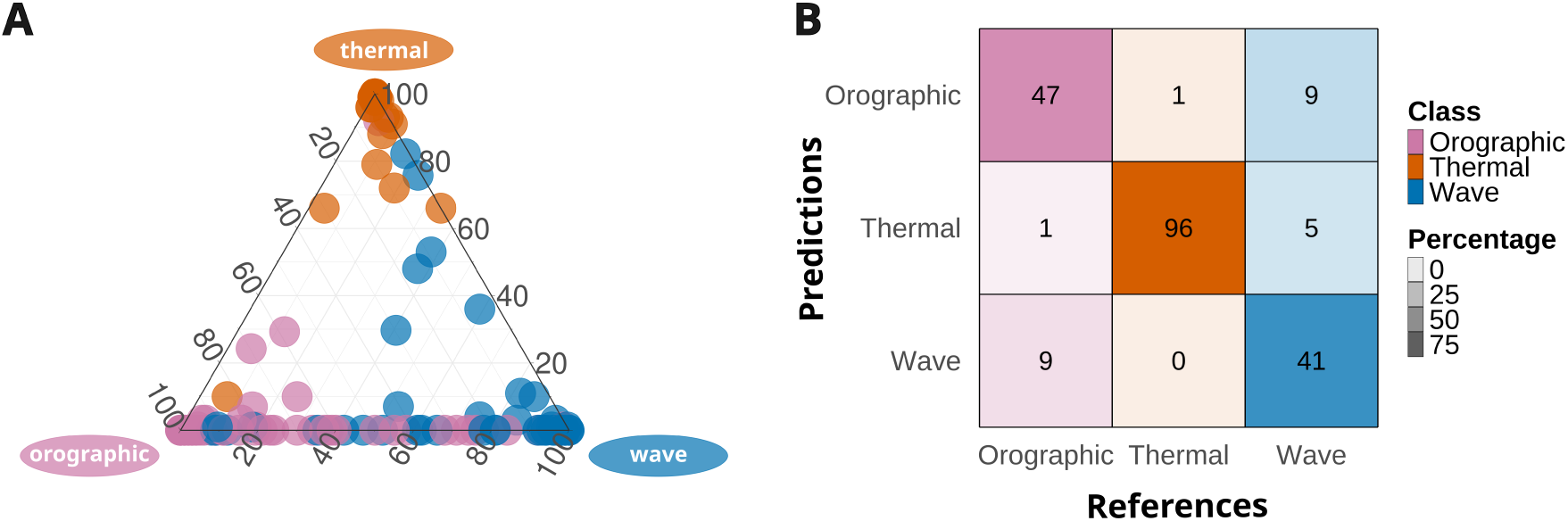
DFA models output (pooled predictions from ten iterations) using environmental predictors: the ternary plot (panel A) and confusion matrix (panel B) show the correct classification of 47/57 orographic uplifts, 96/97 thermal uplifts, and 41/55 gravity waves. The overall accuracy of the ten iterations is 0.88, while the intra-classes accuracies are respectively 0.82, 0.99 and 0.75.

The high accuracy of this model allowed us to predict the uplift sources most likely used during the 25,161 soaring segments we identified between 2020 and 2023 , and those ”available” at the times and locations corresponding to our background dataset (144,800 random points in 2023). To do this, we retrained the DFA algorithm on the full set of 209 manually labelled events and applied this final model to the full dataset of used and available uplifts. The used and available uplift sources predicted by the DFA algorithm were considered reliable only when classified with *>*80% accuracy. For the used dataset, this resulted in 1019 orographic uplifts, 17,167 thermal uplifts and 849 gravity waves for a total of 19,035 accurately classified uplifts, in addition to 823 segments associated to mixed sources (318 orographic/wave, 183 thermal/orographic, 322 thermal/wave) and 5303 to unknown sources. For the available dataset (background points) this resulted in 13,783 orographic uplifts, 78,238 thermal uplifts and 6940 gravity waves for a total of 98,961 accurately classified uplifts, in addition to 8278 mixed sources (4767 orographic/wave, 1528 thermal/orographic, 1983 thermal/wave) and 37,561 unknown sources. The reliability of the accurately classified uplifts used was further validated by comparing the distribution of the weather predictor variables between these automatically labelled sources and the 209 manually labelled soaring segments (see SM section S3 and Figures S1-S2-S3).

Finally, in order to compare the class-specific distribution of the 19,035 uplifts used and the 98,961 uplifts available throughout the year, we used only the data from 2023 (during which data were available continuously without interruptions), for a total of 15,367 segments. For simplicity, and following Carrard et al. (2025), we re-grouped the three uplift sources in two categories, based on their origin: convective (thermals) and mechanic (orographic uplifts and gravity waves, hereafter named together after dynamic uplifts). We then visually compared the distribution of used and available uplifts of these two categories in the different months of 2023.

### Comparison of flight behaviour between uplift sources

We assumed that if eagles’ adopted a different behaviour when using different uplift sources, a classification algorithm run using only behavioural metrics as predictors should return a similar classification of uplift sources to that obtained using environmental parameters only. To test this, we used the predicted classification of the 19,035 soaring segments (only the ”used” uplifts) produced in the previous step. The large size of this dataset allowed us to use a random forest classification algorithm (Breiman 2001, Liaw & Wiener 2002), which is well suited for high-dimensional data and complex, non-linear relationships. This time, we used the uplift classes predicted by the DFA algorithm in the previous step as response variable and the behavioural metrics derived from the biologging (GPS-IMU) data as predictors. Given the high number of behavioural variables calculated from GPS and IMU data (59 in total), we performed a Principal Component Analysis (PCA) (Lê et al. 2008, Kassambara & Mundt 2020) to reduce the dataset dimensionality. We retained the first 10 principal components (PCs), which together explained 79.01% of the total variance. As for the DFA models, also for the random forest we applied a stratified 10-fold cross-validation, balanced for uplift types. Across the ten iterations, one fold was held out as test dataset (10% of the data, on average 2000 segments) while the other nine folds (90% of the data) were used as training dataset. Because of the much higher representation, in the training dataset, of thermal segments relative to orographic and gravity wave segments, we applied a class-balanced random forest (Chen et al. 2004): each bootstrap tree was trained on an equal number of cases per uplift source, drawn independently for every tree from the respective class pools. This approach prevented the disproportionately large thermal sample from dominating model fitting. As for the DFA models, the random forest predictions were pooled across all ten iterations into an aggregated confusion matrix, in order to evaluate the model in terms of overall accuracy (ratio between correctly classified and total uplifts), intra-class accuracy (ratio between correctly classified and total uplifts for each uplift class) and effective accuracy (standard AUC for each class and multi-class AUC as defined by Hand & Till (2001)). Additionally, we derived the receiver operating characteristic (ROC) curve for each uplift class to visualize the true and false positive rates.

### Comparison of COSMO and eagle-derived vertical wind velocities

Vertical wind velocity was extracted from COSMO and annotated to the soaring segments as part of the previous analytical steps. Based on the literature, this same weather parameter can be derived from eagle’s GPS data under specific theoretical assumptions, by subtracting the bird’s sinking speed relative to the air from its observed climbing rate: the residual between what the bird would sink at in still air and how fast it is climbing reflects the upward motion of the air carrying it. While climbing rate is directly measured by the GPS with a known error, the sinking speed cannot be directly measured because it depends on the bird’s morphology and the motion of the air, which itself is modelled. We therefore followed Treep et al. (2016) in estimating the bird’s sinking speed using the Pennycuick aerodynamic model (Pennycuick 1971); we parameterised it with a range of plausible combination of morphologies (body mass, wing span and wing area) for this species and bank angle taken from the literature (see below); and we used COSMO horizontal wind speed as the closest approximation of real air motion in the airspeed calculation. We therefore accept that the resulting vertical wind velocity estimate combines a directly measured climb rate with a modelled sink rate and should be interpreted as an approximation and proof of concept rather than a direct measurement. This approximation is only physically valid during circular soaring as during circular soaring birds actively minimise their sink rate relative to the air, and the Pennycuick (1971) sinking speed equation includes a bank angle correction that is only physically meaningful under steady banked flight conditions. We therefore first isolated circular soaring segments by relying on circling rates published in the literature. Data from Garstang et al. (2022) show that a golden eagle in a gravity wave can complete a full 360 degrees circle in 20-22 seconds (that is a circling rate of 18-16 degrees per second). Other soaring species such as storks, are known to perform a complete soaring circle in a thermal in about 18 seconds (Weinzierl et al. 2016), hence with an average circling speed of 20 degrees per second. We chose a minimum circling rate threshold of 20 degrees per second, to select smaller and less turbulent circling conditions. We thus obtained 5899 climbing segments of circular soaring recorded for all 24 individuals in 2020 and 2023, which we considered suitable to derive vertical wind velocity, by subtracting the birds sinking speed:

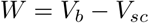

where *W* is the vertical wind velocity, *V_b_*is the bird’s climbing rate and *V_sc_* is the bird’s sinking speed while circling, which was approximated knowing the species’ wing morphology and airspeed, following (Pennycuick 1971):

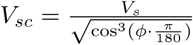

where Φ is the eagle’s bank angle (degrees) and *V_s_* is the sink rate of a soaring bird in relation to the air (m s*^−^*^1^). For simplicity, we assumed a bank angle of 25 degrees as in Garstang et al. (2022), and drag coefficients are from Pennycuick (2008). The sink rate was calculated as follows:

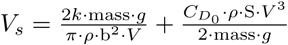

where *g* and *ρ* are gravitational acceleration (m s*^−^*^2^) and air density (at sea level, kg m*^−^*^3^) respectively (both assumed constant); *k* and *C_D_*_0_ are species-specific coefficient (the drag coefficient related to the wings’ efficiency in producing lift, and the zero-lift drag coefficients related to bird’s size and shape respectively); *mass* is the birds’ body mass (kg), *b* is the wing span (m), *S* is the wing area (m^2^) , and *V* is the bird’s airspeed (m s*^−^*^1^). Birds wing area, wing span and body mass were taken from a range of possible values for golden eagles, derived from the literature (Edited by Billerman et al. 2022). To obtain more realistic results, we calculated 106 plausible combinations of body mass, wing span and area, for small, medium and large golden eagles (see SM section S7 Table 1 for the computed constant values). Airspeed *V* was calculated, for each segment, based on the COSMO wind data and the eagles’ ground speed, as follows:

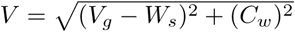

where *V_g_* is the eagles’ ground speed (m s*^−^*^1^), *W_s_* is the wind support (m s*^−^*^1^) and *C_w_* is the cross-wind (m s*^−^*^1^). Wind support and cross-wind were calculated as in Safi et al. (2013).

We visually compared the 5899 bird-derived vertical wind speed values with the values derived from COSMO at those same locations, differentiating for the three uplift types. For 4448 out of the 5899 circular segments, the uplift sources powering them could be accurately classified based on environmental predictors (specific prediction probability *>* 0.8, see DFA model). For this subset of segments, the comparison between COSMO and eagle-derived vertical wind velocities could be separated by uplift source.

## 4 Results

### Automated classification of uplift sources based on environmental parameters

Using atmospheric and topographic proxies, we were able to classify with high accuracy (DFA minimum and maximum accuracy over the 10-fold cross-validation on the test dataset: 0.76 - 1) the uplift sources used by the eagles as well as those available to them at the time and in the area they were flying. Intra-class accuracies were respectively 0.67 to 1 for orographic uplifts, 0.90 to 1 for thermal uplifts, 0.40 to 1 for gravity waves (Figure 1, see also corresponding paragraph in Methods). Applying this model to the entire dataset of 25,161 soaring segments (used uplifts) led to the accurate classification (specific prediction probability *>* 0.8) of 76% of the uplift sources associated to these events (i.e. 19,035 events, of which 17,167 thermals, 1019 orographic uplifts and 849 gravity waves). The reliability of this classification was further confirmed by the comparison of the weather parameters characterising each source in the automatically and manually labelled datasets, both showing a distribution of parameters consistent with the known physical properties of each source (higher orographic lifting coefficient for orographic uplifts, lower sensible heat flux in thermals and higher Brunt–Väisälä frequency in waves) (see SM section S3 and Figures S1-S2-S3). These accurately classified segments constituted the training dataset used in the following analytical step, comparing the flight behaviour of eagles between sources (see corresponding paragraph below).

Applying this same model to the dataset of available sources let to the accurate classification of 98,961 uplifts (of which 3,783 orographic uplifts, 78,238 thermals and 940 gravity waves). The comparison of used vs available uplift sources throughout the year 2023 showed an opportunistic behaviour of golden eagles (Figure 2). In the summer months, when extended daylight hours and increased surface heating favour convection, eagles relied mostly on thermal uplifts. In the colder winter months instead, eagles relied heavily on dynamic uplifts (constituting up to 85.26% of the used uplifts in November) following their seasonal availability, as these are generated in windy conditions independently of solar radiation (Pirotta et al. 2018). However, it is clear that eagles were able to locate and use thermals also in winter, despite their general seasonal scarcity, suggesting a general preference of eagles for this uplift source that aligns with existing literature based on field observations (Kerlinger 1989). It is to be noted that the larger sample size (in terms of absolute number) of recorded soaring events during summer months can be attributed to the tags’ ability to capture more high-resolution data bursts due to fully charged solar batteries (Figure 2B).

**Figure 2:**
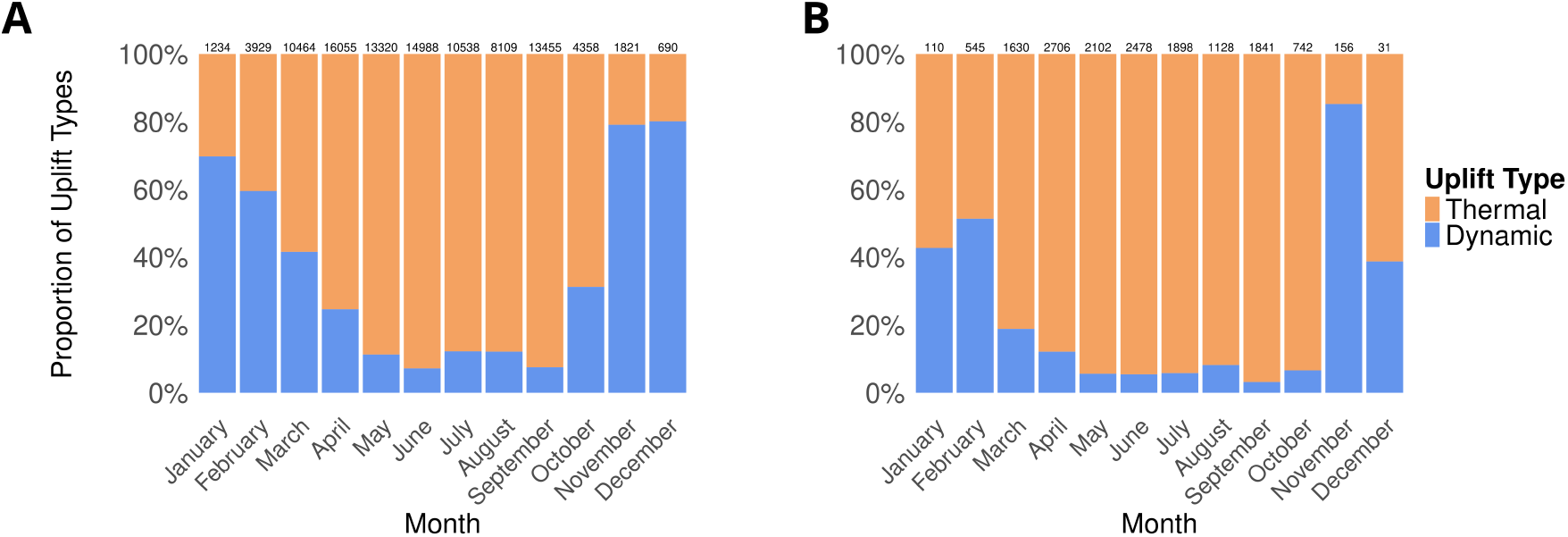
Monthly distribution of uplift sources available (panel A) and used (panel B) throughout the year 2023. In both panels monthly sample sizes are indicated above the corresponding bar.

In addition to the accurately classified sources constituting the used vs available comparison, the algorithm identified a high number of mixed uplift events in both the used and available datasets: 823 mixed events among the used uplifts (of which 183 thermal/orographic, 322 thermal/wave and 318 orographic/wave) and 8278 among the available uplifts (of which 1528 thermal/orographic, 1983 thermal/wave and 4767 orographic/wave). These mixed categories indicate that some events are characterised by intermediate values in the proxies used that do not allow the statistical model to disentangle them, probably due to the co-occurrence, in specific cases, of both mechanic- and convective-driven air flows.

#### Comparison of flight behaviour between uplift sources

Eagles’ flight behaviour was highly diverse within the same soaring segment, showing a behavioural gradient ranging between linear and circular soaring associated to a single uplift (Figure S4) and a high overlap in most behavioural metrics between uplift sources (Figure 3, SM section S7 Table 2). In a single soaring bout, the number of circles performed varied greatly in all three types of uplift sources, ranging from 0 to 29.54 in orographic uplifts, 0 to 61.07 in thermal updrafts and 0 to 33.63 in gravity waves. The comparison of behavioural flight parameters across the three uplift sources revealed striking similarities in the large majority of behavioural metrics, with means differing by less than 15% and standard deviations consistently large relative to between-source differences (SM section S7 Table 2). The most notable distinctions were found in height above ground and energetic proxies: gravity waves were associated with flight at greater altitude and higher values of VeDBA and acceleration variance on the vertical axis, suggesting more energetically demanding flight in more turbulent conditions, while thermals were associated with a higher number of circles per soaring segment, consistent with the compact spatial extent of convective uplifts. All other parameters — including ground speed, turning angle, circling rate, body tilt and Euler angles — showed broadly overlapping distributions across uplift sources (Figure 3), providing a first indication that eagles do not adopt systematically distinct flight strategies in response to different uplift types.

**Figure 3:**
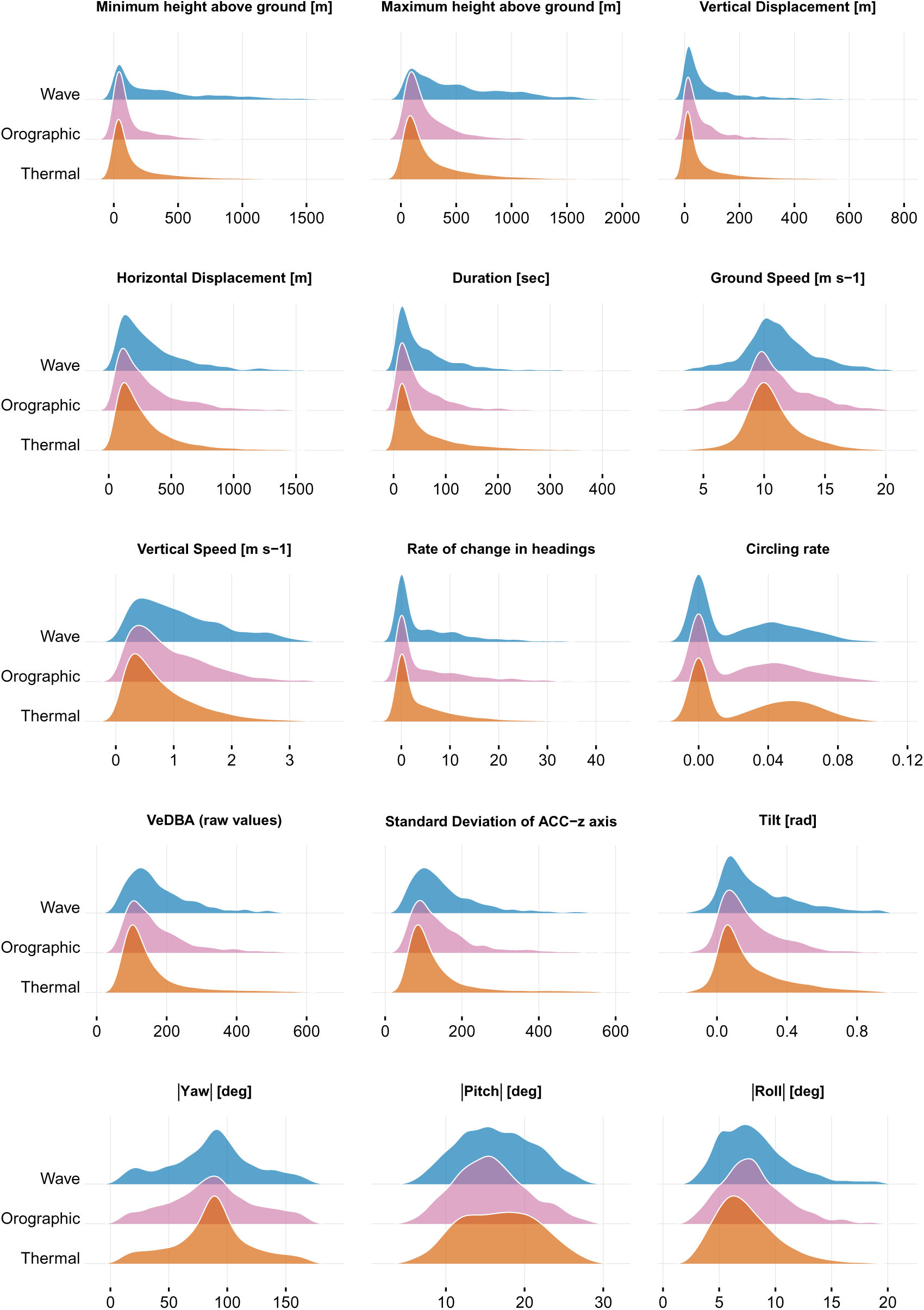
Density distributions of soaring flight variables by uplift type, averaged along soaring segments. For visualization purposes, the x-axis was restricted to the central 99% of the data (excluding the lowest and highest 0.5%).

The first 10 principal components (PCs) explained 79.01% of the variance in the behavioural dataset (Figure S5). The variables that contributed the most to the 10 PCs are metrics relative to: yaw, pitch and roll angles, ODBA and VeDBA, number of circles, turning angle, vertical speed, turbulence (approximated by the average standard deviation of acceleration on z-axis), and vertical displacement.

From the random forest model including the 10 components of the PCA on the behavioural variables, we obtained a consistent overall accuracy ranging between 0.6758 to 0.725 over the ten iterations (Figure S6). Within-class performance varied considerably between classes: the model could correctly classify on average 73.4% of the thermal uplifts, 37,3% of orographic uplift and 43.9% of mountain gravity waves. The predictions of both orographic uplifts and gravity waves showed a higher false negative rate compared to thermal uplifts, and were wrongly attributed, in most cases, to thermals. This is reflected in the multiclass AUC, which has an average value of 0.653. This suggests that the model is not effectively distinguishing between the classes, and hence that, at least for this species, behaviour alone is not sufficient to determine the uplift source that eagles are flying in. To account for the full variability of behaviour within uplift sources, we trained a second random forest with the original frequency-weighted uplift classes without subsampling and balancing the relative proportions of uplift classes, but the classification power of the model did not improve (see SM section S6).

#### Comparison of COSMO and eagle-derived vertical wind velocities

We visually compared the density distribution of the vertical wind velocity extracted from COSMO with that derived from the eagles’ climbing rate during circular soaring. We removed outliers (unrealistically low and high values) by excluding the first 0.05% and last 0.05% of the data. The vertical wind velocity extracted from COSMO exhibits a density distribution centred on 0 m/s, whereas the one derived from the eagles’ climbing rate shows a clear shift towards higher velocities, with an average value of 4.27 m/s (Figure 4A). Considering that eagles are effectively climbing the air column, the vertical wind speed values derived from their flight patterns appear to be more biologically realistic relative to their relative high wing loading (7.2 kg/m^2^, Bohrer et al. (2012)). COSMO velocity values were clustered around 0 m/s for thermal uplifts, shifted towards positive values in orographic uplifts, and towards negative values in gravity waves (Figure 4B). In contrast, the vertical wind speed extracted from the eagles’ flight demonstrates a dense distribution towards the same range of positive values across all uplift types: a more realistic representation of the positive component that must have been available at fine scales, given the physical constraints associated to soaring flight. Despite both measurements not being accurate representation of the true vertical wind velocity at that time and location, this comparison highlights that even kilometre-scale weather models are insufficient to represent atmospheric conditions experienced by soaring birds. For thermals, this discrepancy is largely due to the fact that they mostly occur on spatial and temporal scales that cannot be resolved by COSMO. For gravity waves, this could additionally indicate that COSMO correctly models mountain waves, but fails to capture their spatial distribution and the strength of the associated vertical velocity field. This spatial mismatch could explain why gravity waves segments are associated with negative vertical velocities on average (Figure 4B). Orographic uplifts are more consistent between the two estimates, which reflects the fact that a positive orographic uplift coefficient (i.e. positive vertical velocities) was used as a criterion to attribute a soaring segment to this category.

**Figure 4:**
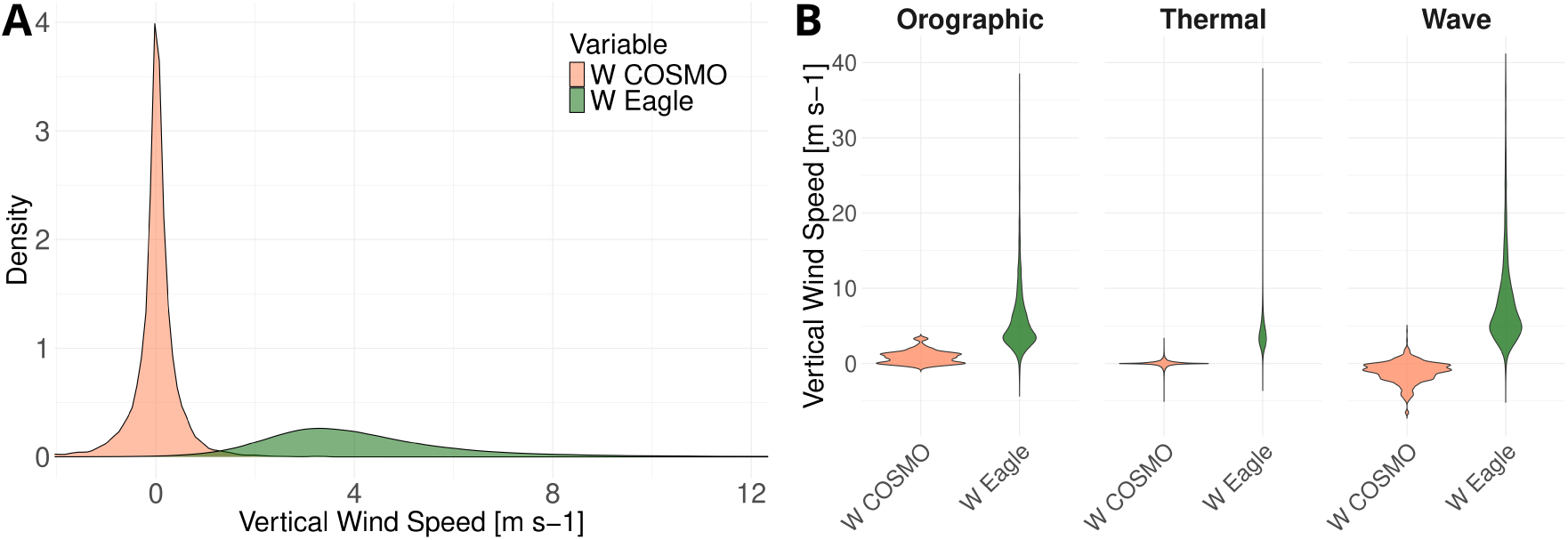
Validation of eagle-borne vertical wind speed. Panel A: Density distribution of vertical wind speed estimated by COSMO and eagles. For visualization purposes, the x-axis was limited to the 1st–99th percentiles. Panel B: Density distribution of vertical wind speed estimated by COSMO and eagles, shown separately for each uplift source.

## 5 Discussion

By combining high-resolution movement and weather data, we offer novel insights into the soaring flexibility of golden eagles in topographically complex regions, revealing at unprecedented resolution not only how birds respond to atmospheric dynamics, but also how their behaviour could be used to characterise them. Using high resolution weather proxies extracted from the regional weather model COSMO, we developed a first automated algorithm for the classification of uplift sources into thermals, orographic uplifts and gravity waves. This allowed us to accurately classify the uplift sources used by the eagles to support 19,035 soaring events in 2020 and 2023. We quantified, for the first time, the contribution of mountain gravity waves to golden eagles’ soaring behaviour, after their use had been suggested in the late 90s by Kerlinger (1989), then anecdotally shown by Garstang et al. (2022) on one golden eagle in the Appalachian, and more recently demonstrated across several golden eagles in the Alps by Carrard et al. (2025). The same algorithm independently identified soaring events powered by more than one uplift source simultaneously (mixed uplift sources). Our comparison of uplift sources used (and unequivocally classified) with their availability throughout the year suggests that eagles’ make opportunistic use of uplift sources available seasonally, with the distribution of uplift sources used broadly reflecting their availability in the environment. The analysis of the behavioural metrics associated to these same soaring events suggested that eagles could flexibly use different uplift sources without the need to adapt their flight behaviour. This indicates that behaviour alone cannot inform us about the specific source of uplift used, and possibly, that such a strict classification of uplift sources might not be representative of real atmospheric dynamics. Together with the large number of mixed uplift events identified by our algorithm, these results suggest that both atmospheric conditions and behavioural responses may be better described as continua than as discrete categories. Finally, our comparison of vertical wind velocities extracted from COSMO and derived from the eagles’ climbing rates highlights that kilometre-scale weather models are still insufficient to represent atmospheric conditions relevant for soaring birds. While the observed discrepancy mostly reflects the fact that uplifts largely occur on scales unresolved by COSMO, the important mismatch observed for dynamically induced updrafts (mostly for gravity waves) also suggest that the model misrepresents longer-lived atmospheric patterns such as windward orographic lifting and mountain gravity waves. While this remains to be investigated in more details, this latter observation suggests that outputs from numerical weather models could benefit from the integration of movement-derived measurements.

### Opportunistic seasonal use of uplift sources

Our classification of uplift sources based on high-resolution atmospheric data allowed us to determine with unprecedented clarity the uplift sources powering the soaring flight of 24 eagles throughout the year. To our knowledge, this is the first in depth quantification of used versus available uplift sources and it allowed us to substantiate seasonal patterns of eagles’ uplift use already proposed in the literature (Pirotta et al. 2018, Murgatroyd et al. 2018). In fact, the predominant use of thermal uplifts by golden eagles had already been suggested based on a combination of observational data, behavioural data and synoptic weather conditions (Kerlinger 1989, Duerr et al. 2012, 2015, Katzner et al. 2015, Bohrer et al. 2012). Our approach decouples uplift identification from behavioural observation for the first time, and confirms this pattern. Thermals represented the main uplift source for eagles throughout the year but during winter months, when thermals became less available, eagles flexibly relied on dynamic uplift sources, which contributed to a minimum of 3.2% to a maximum 85.3% of the soaring events. Importantly, the monthly contribution of mountain gravity waves, quantified here for the first time, ranges between 0.8% to as high as 74.3%.

#### Behavioural flexibility across uplift sources

The intrinsic differences between thermal and dynamic uplifts in their spatio-temporal distribution, strength, stability, and potential for altitude gain (Organisation Scientifique et Technique Internationale du Vol à Voile 2009, Shamoun-Baranes et al. 2003) led us to expect that golden eagles would adjust their soaring behaviour to these properties, and therefore that distinct flight patterns could help us identify the source of uplift even in the absence of atmospheric information. Instead, the distribution of all behavioural parameters largely overlapped between uplift sources, with means differing by less than 15% and standard deviations consistently larger relative to between-source differences. This lack of behavioural differentiation was qualitatively supported by our visual inspection of the flight trajectories, showing similar fight patterns across all uplift sources, and quantitatively reflected in the performance of our machine learning classifier, which failed to effectively distinguish between the three sources when trained on behavioural parameters alone. This finding challenges the traditional view that associates binary flight modes with particular uplift types: while linear soaring is generally linked to orographic uplifts and circular soaring to thermals (Alerstam & Hedenström 1998, Kerlinger 1989), our results reveal that eagles use both flight modes across all three uplift sources. From the ground speed to the circling rate, this consistent pattern suggests a highly versatile flight strategy that allows golden eagles to efficiently exploit a wide range of environmental conditions, without requiring significant alterations in their soaring behaviour. This flexibility may reflect a highly adaptive generalised soaring strategy, shaped by the selective advantage of maintaining efficient flight across the full range of atmospheric conditions encountered in a topographically complex landscape. Eagles adaptability also owes to their morphology and maneuverability, and future exploration of similar behavioural parameters in species with higher wing loading, e.g. in large vulture species, could result in stronger behavioural differences between uplift sources.

#### Atmospheric processes and behaviour interact along a continuum

From an evolutionary perspective, the versatility of soaring behaviour might also be a response to the complex and often mixed nature of atmospheric dynamics in the boundary layer, where convection and mechanically-induced airflows can contribute to the same uplift powering a single soaring event, as recently suggested also by Carrard et al. (2025). This is particularly true over complex terrain, where the combination of synoptic airflow, topography, and regional thermal effects can lead to the superposition of convective and dynamically induced updrafts. This phenomenon, which we are only starting to be able to investigate thanks to the higher resolution of weather models, questions the need for such strict categorisation of both behaviour and uplift sources often suggested in the literature (Alerstam & Lindström 1990, Kerlinger 1989, Murgatroyd et al. 2018, Santos et al. 2017) but probably not realistic. The algorithm developed in our study for the automated classification of uplift sources was trained on events that could be unambiguously labelled as one of three categories via manual inspection in the recent study by Carrard et al. (2025). Yet, when applied to our full dataset, this algorithm independently identified 3.3% and 5.7% of the uplifts analysed (used and available respectively) as mixed, i.e. powered by more than one uplift source, with each contributing at least 0.4 of probability. This finding confirms that strict categorical classification of uplift sources, while analytically tractable, does not reflect the physical reality of the atmospheric boundary layer, which is in line with results obtained through visual inspection by Carrard et al. (2025). Therefore, the amount of mixed thermal-dynamic events in our dataset and the flexibility in the eagles’ behaviours provide clear indications that both atmospheric conditions and behavioural responses exist along a continuum and vary within each uplift event.

#### Soaring flight highlights limitations and opportunities for high-resolution weather modelling

Despite regional weather models like COSMO provide new insights into complex atmospheric phenomena such as gravity waves, local orographic lifting, and deep convection, the model resolution remains insufficient to fully capture the conditions experienced by soaring birds, as illustrated by the discrepancies in wind vertical velocities estimated by COSMO and derived form the eagles’ climbing rate. This resolution gap also has implications for regional weather forecasting, as it contributes to substantial uncertainties in the prediction of storm timing and intensity (Sanders & Doswell III 1995, Mandement & Caumont 2020). The overall comparison between vertical velocities across uplift sources clearly illustrates this resolution gap. However, while the discrepancy associated to thermal uplifts can be largely attributed to the too coarse resolution of COSMO (in both space and time) relative to the duration and extent of thermals, the large mismatch observed for gravity waves, which are instead often resolved by COSMO (Carrard et al. 2025), suggests that mountain waves are present but often inaccurately represented in the model. In the future, the use of flight data could potentially be used to address both sources of this discrepancy. On the one hand, as a ground truth to test the accuracy of regional numerical weather model and possibly improve the current representation of dynamically induced updrafts and deep convection over complex terrain. On the other hand, eagles’ climbing rates could be used to test the ability of the model to parametrise short-lived thermals and dynamically induced turbulent uplifts, that are currently unresolved but that are shown to sustain soaring flight, and potentially improve these parametrisations.

Testing and comparing the accuracy of either COSMO and eagle-derived measurements was not possible, as this would require a three-dimensional anemometer measuring the same air column where the eagle is flying. However, while both are not direct measures of this parameter but the result of models and equations under specific assumptions, their comparison highlights a real gap in our current representation of atmospheric dynamics, as well as opportunities to bridge it. Next generation decametre-scale weather models hold promises in this direction (Krieger et al. 2025), but their parametrisation also relies on direct observations of complex airflows and atmospheric parameters that are challenging to measure, and lately targeted by specific measurement campaigns (Ward & Westerhuis 2023, Bugnard et al. 2025). Soaring birds, during their daily movements, routinely sample these same airflows that such campaigns are targeting. The results of our comparison therefore support the growing view that animal movement patterns encode precious environmental information that can expand the current environment observation network and inform the parametrisation of weather models (Ellis-Soto et al. 2023) to potentially increase their accuracy, as already the case for oceanographic models (McMahon et al. 2021). Building on previous studies on bird-derived atmospheric measurements (Weinzierl et al. 2016, Goto et al. 2017, Yonehara et al. 2016, Treep et al. 2016), which relied on coarser atmospheric products for comparison or short time-windows, the results of our comparison against the kilometre-scale COSMO model advocates for the potential of biologging-derived meteorological variables, even if direct validation remains elusive. Future studies should aim at directly validating these variables against measurements from traditional weather sensors, and test the improvement that the integration of these data could achieve. Finally, more in-depth analysis of GPS and IMU data encoding point-by-point bird posture might offer further insights into the fine-scale adjustments soaring birds make in response to differently turbulent uplift sources, potentially increasing the level of environmental detail these behavioural data could resolve (Williams et al. 2015).

## 6 Conclusions

This study illustrates both the promise and the remaining challenges of closing the resolution gap between animal behaviour and environmental data. For decades, the inability to characterise uplifts at the scale at which birds experience them forced researchers to infer atmospheric context from behavioural proxies and coarse synoptic conditions; a circularity that this study begins to break. Coupling sub-second biologging with kilometre-scale weather modelling opens a new observational window simultaneously in two directions: inward, toward a more mechanistic understanding of how individual animals perceive, respond to, and make decisions within dynamic physical environments; and outward, toward a finer characterisation of those environments themselves, in places and at scales where conventional sensor networks remain sparse. As biologging, remote sensing and atmospheric modelling continue to converge in resolution, this bidirectional exchange will become increasingly tractable, and the interface between behavioural ecology and environmental science increasingly productive. A key challenge going forward will be to determine how increased environmental and behavioural realism translates across spatial and temporal scales, that is, from individual soaring decisions to migratory routes and species distributions. More broadly, our results suggest that some long-standing assumptions about the specificity of behavioural responses to environmental conditions may reflect the limits of what could previously be measured rather than true biological constraints on how animals respond to their physical environment. Revising these assumptions will require not only finer-scale measurements, but also analytical frameworks capable of embracing the continuous nature of both atmospheric processes and behavioural responses.

## Acknowledgments

We thank MeteoSwiss and particularly Lukas Jansing for providing the COSMO-1 analysis data. We are grateful to David Jenny and Enrico Bassi for their golden eagles’ data contribution; Adriano Greco, Alessandro Mercogliano, Andrea Roverselli and Klaus Bliem for their support during the eagles’ tagging. We are grateful for the crucial assistance of the mountain rescue teams of the Guardia di Finanza as well as the forestry and game wardens of South Tyrol/Alto Adige and the national park team in Gesäuse, as well as the ERSAF-Stelvio national park Italy and the Department of Wildlife and Fishery Service Grisons, Switzerland (AJF) and many gamekeepers. We thank the accompanying veterinarians Drs Michel Mottini and Gilberto Volcan for their time and expertise in the field. We are also grateful to Emily Shepard and Sergio Lambertucci for insightful feedback on an advanced draft of this manuscript.

## Ethics

The handling and ringing of golden eagle nestlings in Switzerland was carried out under the authorization of the Food Safety and Veterinary Office Grisons, permit nos. GR 2017 06, GR 2018 05E, GR 2019 03E, GR/08/2021, and the Federal Office for the Environment, licence no. TV201903E. In Italy, the permissions for handling, tagging and marking were obtained from the autonomous region of South Tyrol (Dekret 12257/2018 and Dekret 8788/2020), as well as from the region of Lombardia for ringing and tagging through in Lombardia and South Tyrol by ISPRA (Istituto Superiore per la Protezione e la Ricerca Ambientale) with the Richiesta di autorizzazione alla cattura di fauna selvatica per scopi scientifici (l.r. 26/93). In Austria, all procedures for handling, tagging and marking were approved by the Ethics Committee of the University of Vienna (no. 2020-008) and permitted by the Federal Ministry for Education, Science and Research (no. 2020-0.547.571), Styria (BHLI-165942/2021-2) and Upper Austria (LFW-2021-263262/7-Sr).

Finally, in Germany birds handled, tagged and ringed were done so under the permission issued by the government of Oberbayern (2532.Vet 02-16-88 and 2532.Vet 02-20-86). All procedures followed the ASAB/ABS guidelines for the ethical treatment of animals in behavioural research and teaching and all applicable international, national, and/or institutional guidelines for the care and use of animals were followed. The handling of birds was performed with maximum care and minimal disturbance to nests and the landscape.

## Data accessibility

The R and Python scripts used to process and analyse the data are publicly available on Github at GoldenEagles COSMO pub.git. The raw data containing the complete movement trajectories is deposited on movebank.org (Study Name ”LifeTrack Golden Eagle Alps”, Movebank study ID 282734839). This dataset will be available upon requests made to the data owners (KS, MW, MUG and PS) for research purposes, provided that the data use will not threaten the study populations.

## Declaration of AI use

AI was used to rephrase specific sentences of the manuscript and to debug R and Python code. All text and code was written by humans and only edited and completed in specific parts with the help of AI technologies.

## Authors’ contributions

FF Data curation, Project Administration, Investigation, Methodology, Software, Validation, Formal analysis, Visualization, Writing – original draft, Writing – review and editing; TC Methodology, Formal analysis, Software, Validation, Writing – review and editing; MUG Data curation, Resources, Writing – review and editing; JSH Data curation, Resources, Writing – review and editing; KS Conceptualization, Data curation, Resources, Writing – review and editing; MASp Resources, Writing – review and editing; MW Data curation, Writing – review and editing; PS Data curation, Resources, Writing – review and editing; MS Conceptualization, Formal analysis, Project Administration, Software, Supervision, Writing – original draft, Writing – review and editing.

All authors gave final approval for publication and agreed to be held accountable for the work performed therein.

## Competing interests

The authors declare no competing interests.

## Funding

No funding has been received for this article.

## Supplementary materials

### S1: Behavioural variables definitions

From every GPS location of a soaring segment, we extracted the following information, that were then summarised per segment:

**–** height of the eagle: the height above ellipsoid is recorded by the tag, and we used it to estimate the height above sea level and above the ground of every eagles’ location. For this we relied on elevation data from a digital elevation model (DEM) openly available for Europe at 30 m spatial resolution (Hengl et al. 2020);

**–** horizontal displacement: horizontal distance between the first and the last point of the segment, in metres;

**–** vertical displacement: vertical distance between the first and the last point of the segment, in metres, calculated as difference of the first and last recorded height above ellipsoid;

**–** average vertical speed: vertical distance divided by time lag between consecutive points, averaged along the segment, in metres per second;

**–** ground speed: horizontal distance divided by time lag between consecutive points, averaged along the segment, in metres per second;

**–** cumulative turning angle along the complete segment (in radiants): sum of the angles between all consecutive locations in a segment;

**–** number of change of headings in a segment: change in angle sign while turning, summing the occurrences per segment;

**–** tilt: arctan function of the ratio between vertical and horizontal displacement;

**–** number of circles: the number of circles performed along the soaring segment, calculated as the cumulative turning angle divided by 6.28 radiants (360 degrees);

**–** circling rate: the number of circles performed divided for the duration of the soaring segment (in seconds).

From the ACC data associated to each GPS point, we extracted:

**–** overall dynamic body acceleration (ODBA): calculated following (Wilson et al. 2020) as the non-vectorial sum of the absolute dynamic acceleration values along the three spatial axes and used as a proxy of energy expenditure and flapping probability in flight;

**–** vectorial dynamic body acceleration (VeDBA): calculated following (Wilson et al. 2020) as the vectorial sum of dynamic acceleration along the three spatial axes and used as a proxy of energy expenditure and flapping probability in flight;

**–** standard deviation of the acceleration over the z-axis (vertical ACC axis), used as a proxy of flight turbulence (Laurent et al. 2021).

From the IMU data (combination of accelerometer and gyroscope) associated to each GPS point, we extracted yaw, pitch and roll angles following (Safi et al. 2025):

**–** yaw angle: left-and-right rotation around the z-axis (vertical axis), from head to tail;

**–** pitch angle: up-and-down rotation around the x-axis (horizontal axis), crossing the bird barycenter;

**–** roll angle: side-to-side rotation around the y-axis (longitudinal axis), from one wingtip to the other.

Due to the different duration of the classified soaring segments (from 7 to 776 seconds), we applied a time component normalization of some metrics (such as cumulative turning angle, vertical and horizontal displacement, number of circles of circular soaring, number of changes in heading, cumulative yaw, pitch and roll) and extracted comparable rates (e.g. ’circling rate’ calculate as number of circles divided by segment duration).

### S2: Derivation of secondary variables (COSMO)

The vertical stability of the atmosphere was expressed with the Brunt-Väisälä frequency:

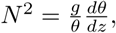

where *g* is the gravitational acceleration, *θ* is the potential temperature, and 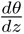 is the vertical gradient of potential temperature with:

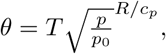

where *T* is the temperature, *p* the atmospheric pressure, *p*_0_ the reference pressure (1000 hPa), *R* the specific gas constant of air and *c_p_* the specific heat capacity at a constant pressure. Both *θ* and *N* ^2^ were calculated for each model level. The maximum stability was taken as the maximum value of *N* ^2^ in the lower 2000 m above ground and the height of this maximum stability was taken as a rough estimate of the height of the boundary layer.

### S3: Validation of automatized uplift classification algorithm

Orographic uplifts are characterized by higher orographic uplift potential (Figure S1 A,B) than thermal uplifts and gravity waves. However, as expected, orographic uplifts and gravity waves are both dynamic uplifts, and therefore show similar distributions for surface sensible heat flux (higher than thermals, Figure S1 C,D), absolute vertical wind speed (higher than thermals, Figure S2 A,B) and horizontal wind speed (higher than thermals, Figure S2 C,D). Gravity waves are identified especially from the Brunt–Väisälä frequency as height of maximum stability (lower than orographic and thermal uplifts, Figure S1 E,F) and value of maximum stability (higher than orographic and thermal uplifts, Figure S1 G,H), which characterize the ideal conditions for mountain gravity waves formation. Thermal uplift are better differentiated from the dynamic uplift sources by a lower value of surface sensible heat flux (Figure S1 C,D), which indicates a thermal convection from the Earth surface towards the atmosphere. The topographic features distribution shows instead a similar pattern across all three uplift types (Figure S3).

**Figure S1:**
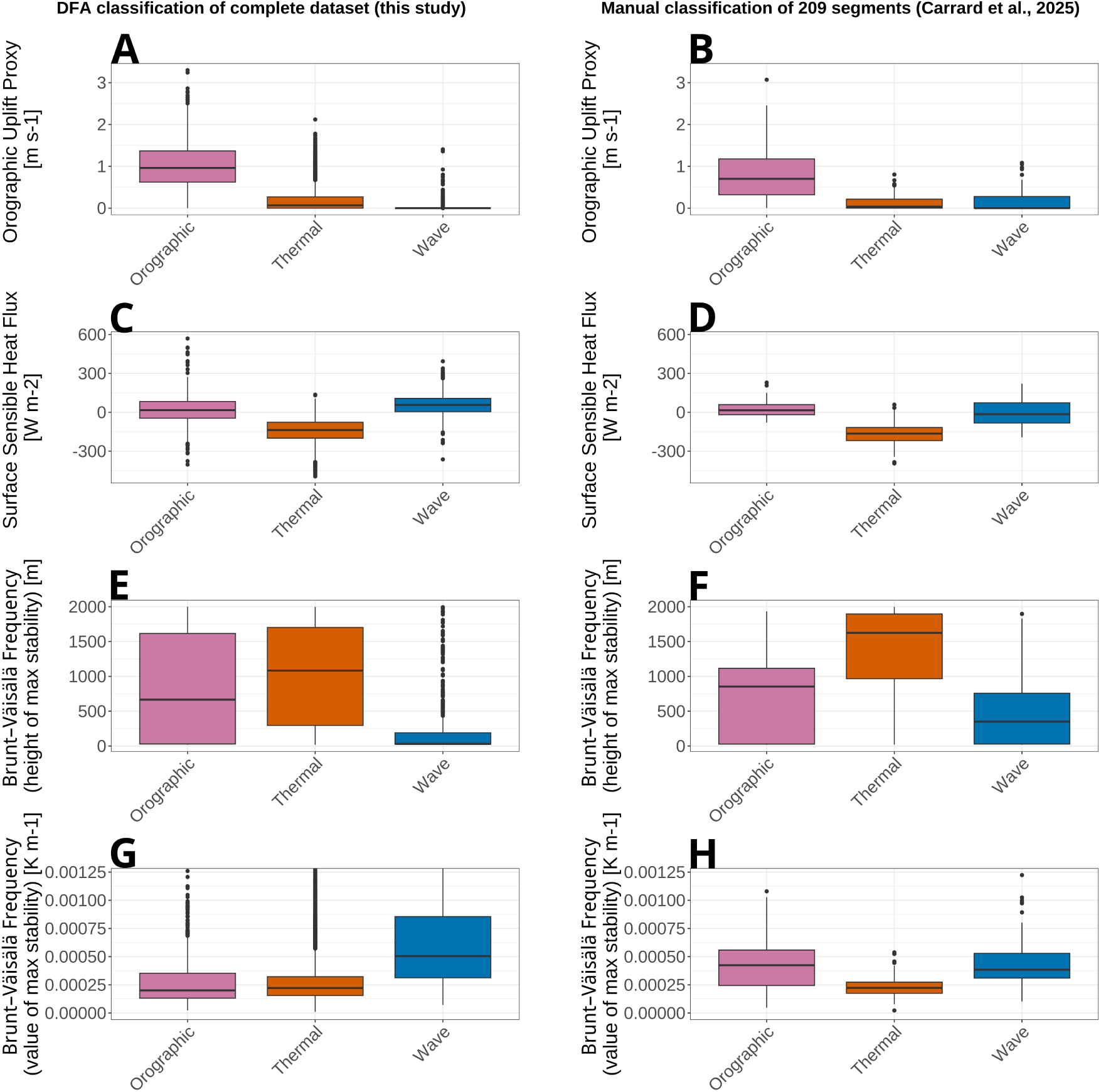
Boxplot distributions of the COSMO atmospheric variables (average values across the soaring segments): orographic uplift proxy (panels A,B) , surface sensible heat flux (C,D), Brunt–Väisälä frequency as height (E,F) and value of maximum stability (G,H). In the right column (panels B,D,F,H) values of the 209 manually classified uplift events. In the left column (panels A,C,E,G) reported values for 19,035 uplift events automatically classified with the DFA algorithm.

**Figure S2:**
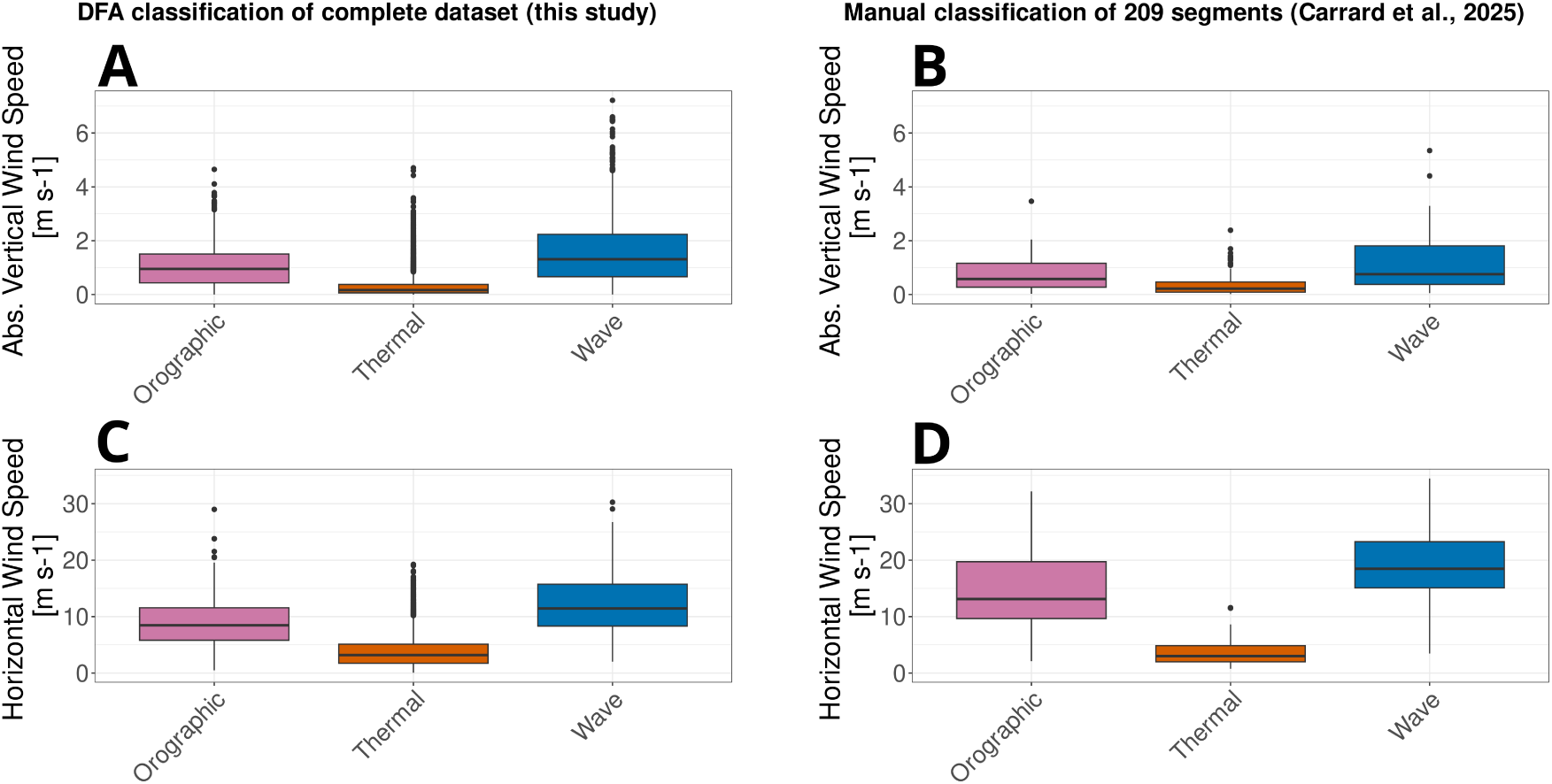
Boxplot distributions of the COSMO wind parameters (average values across the soaring segments): absolute value of vertical wind speed (panels A,B) and horizontal wind speed (C,D). In the right column (panels B, D) values of the 209 manually classified uplift events. In the left column (panels A,C) reported values for 19035 uplift events automatically classified with the DFA algorithm.

**Figure S3:**
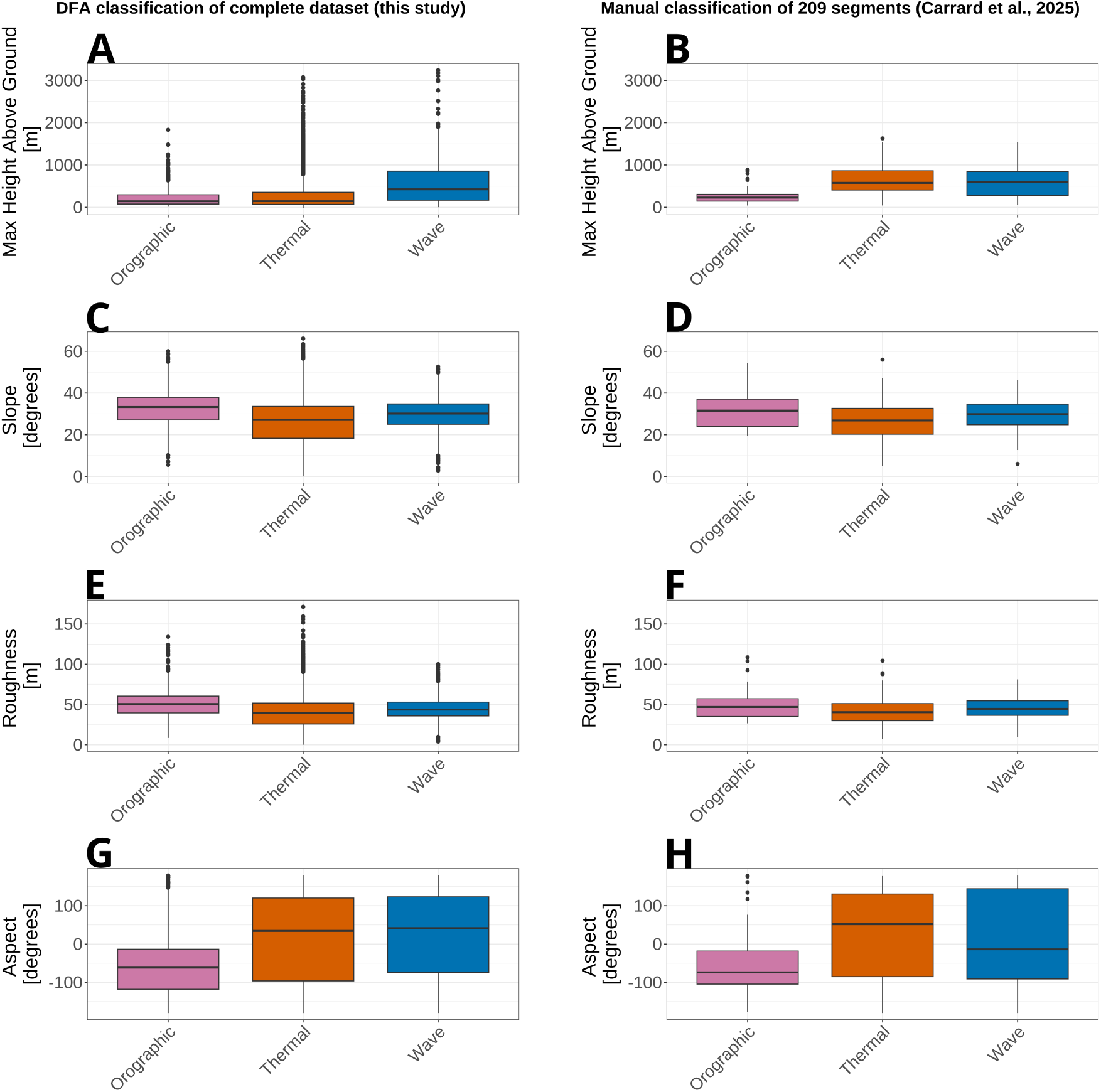
Boxplot distribution of topographic properties (average values across the soaring segments): maximum height above ground (panels A,B), slope (C,D), roughness (E,F), and aspect (G,H). In the right column (panels B,D,F,H) values of the 209 manually classified uplift events. In the left column (panels A,C,E,G) reported values for 19,035 uplift events automatically classified with the DFA algorithm.

### S4: Visualization of trajectories across uplift types

**Figure S4:**
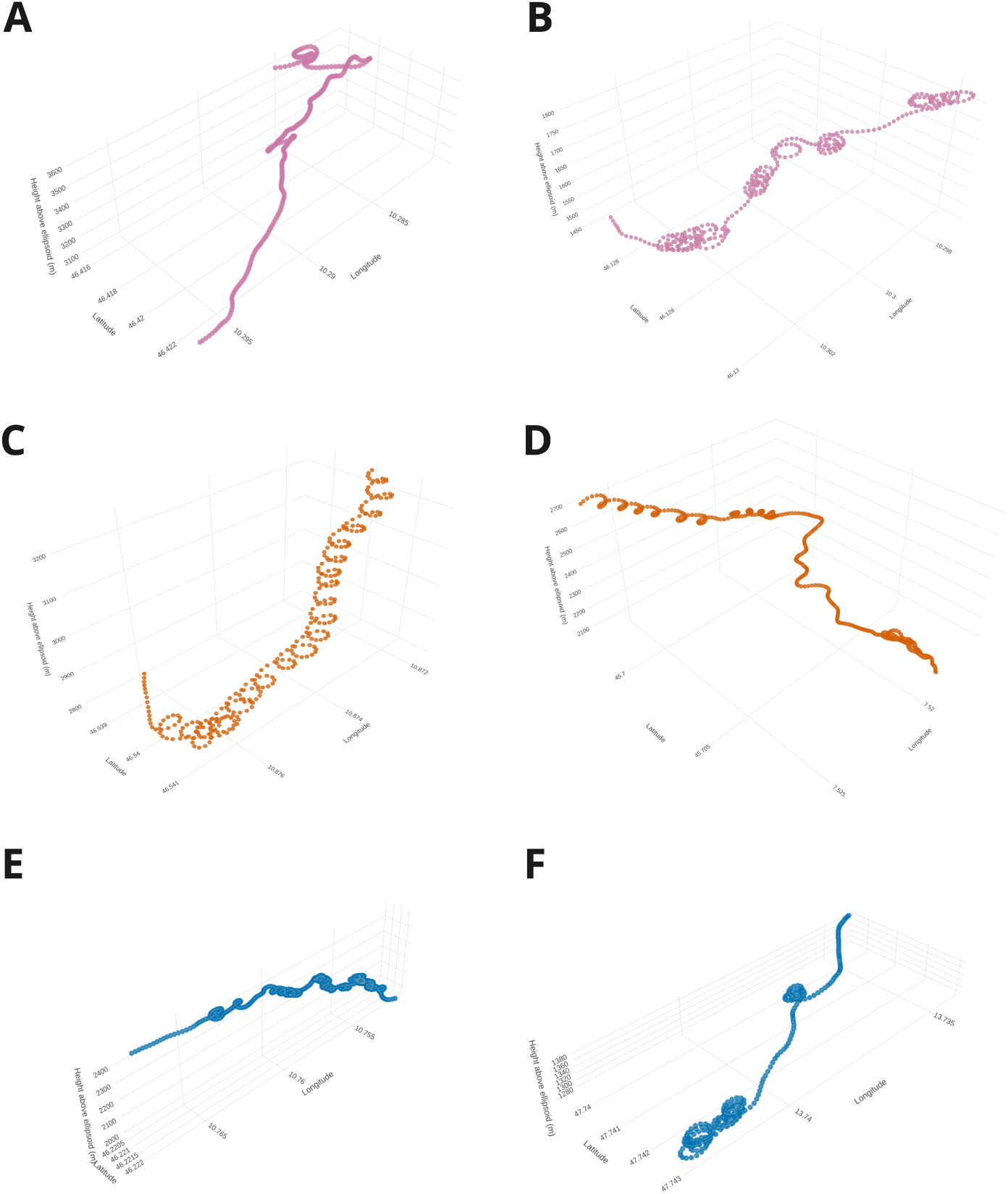
Examples of high-resolution GPS trajectories (1 Hz) of eagles flying in different uplift sources, plotted in 3D over longitude, latitude and height above ellipsoid (in metres). Each row represent a range of flight patterns adopted in each type of uplift source (panels A,B: orographic uplifts; C,D: thermals; E,F: gravity waves), while columns show two examples of flight pattern associated to the same uplift source (left panels A,C: stereotypical behaviour as described in the literature; right panels B,D: typical pattern encountered in our dataset). Previous studies associate orographic uplifts mainly to linear soaring (A) and thermals to circular soaring (C). Behaviour in gravity waves was rarely observed and described. Our data show that all uplift types, including gravity waves, are characterised by a mixture of circular and linear soaring (B, D, E and F).

### S5: PCA on behavioural parameters

**Figure S5:**
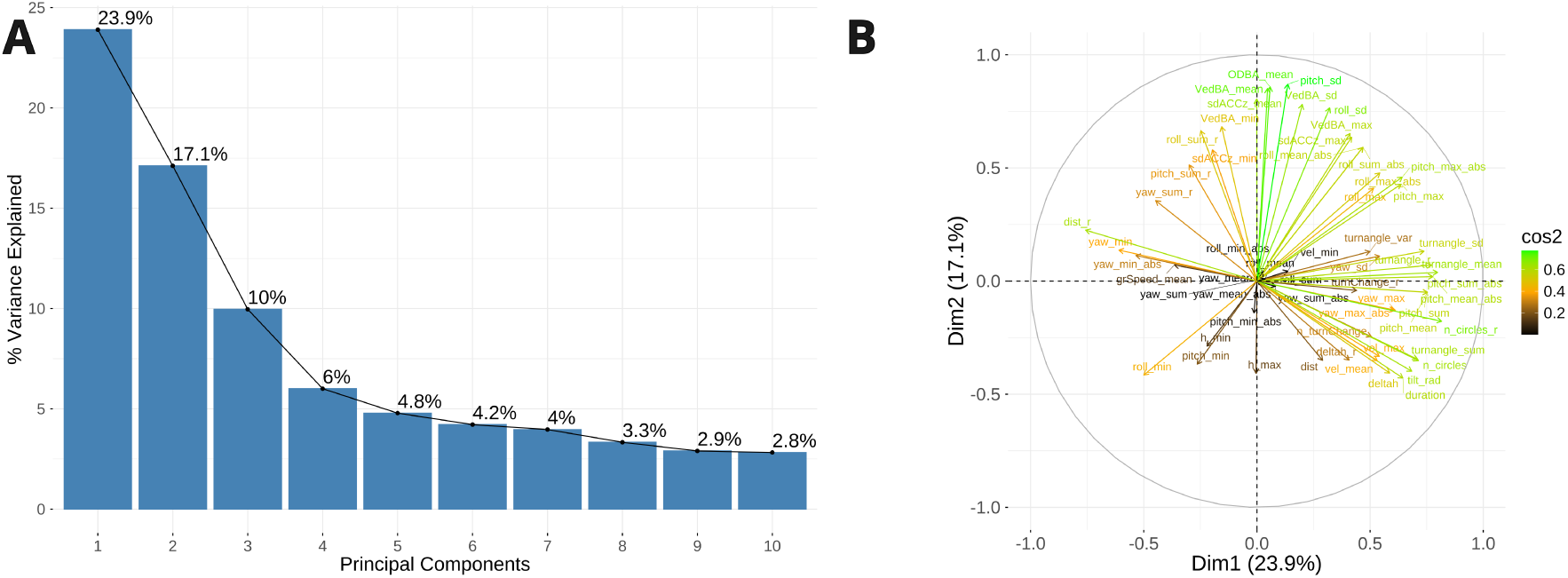
PCA on the behavioural parameters. Panel A: Percentage of variance explained by the first 10 principal components. Panel B: PCA variable plot colored by their quality of representation (cos2 values) for all 59 behavioural variables. Variables are displayed in the space of the first two principal components, coloured by their cos2 value. Variables closer to the circumference and with higher cos2 values suggest better representation of each variable by the first two principal components.

### S6: Random Forest model

**Figure S6:**
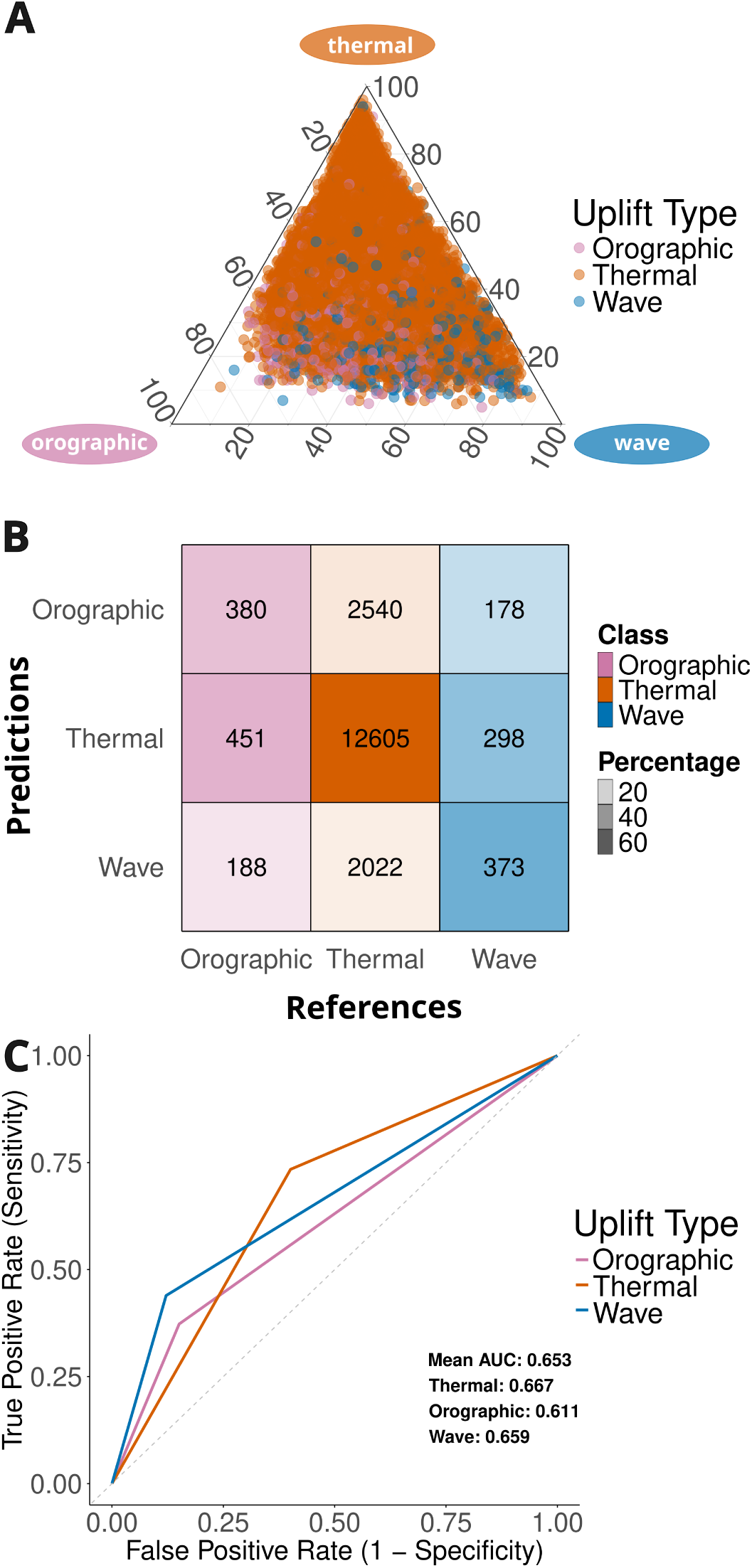
Output of the random forest models (pooled predictions from ten iterations) using behavioural predictors: the ternary plot (panel A) and the confusion matrix (panel B) show the correct classification of 380/1019 orographic uplifts, 12605/17167 thermal uplifts, and 373/849 gravity waves. The overall accuracy of the iterations is 0.702, but the model is incapable of effectively distinguish uplifts from thermal ones. Panel C: ROC curve, multi-class AUC is 0.653.

#### Random Forest with frequency-weighted uplift sources

We trained a second random forest model on the dataset maintaining its original class frequencies, that is, without balancing the relative proportions of uplift types in the training data. We considered this frequency-weighted approach worth testing because it retains the full range of variability within the uplift classes, rather than the equally-sized subsample of the class-balanced approach for each tree of bootstrap. This model therefore reflects both the natural prevalence of uplift types and the full behavioural variance of the dominant class (thermals constitute the 90.2% of all predicted uplift types). We obtained a consistent overall accuracy ranging between 0.8997 and 0.9043 over the ten iterations (Figure S7). Despite the high overall accuracy of the model, its within-class performance varied a lot between classes. While the model could correctly classify on average 99.9% of the thermal uplifts, it could only classify 0.88% and 1.18% of orographic uplift and mountain gravity waves, respectively. The predictions of both orographic uplifts and gravity waves showed in fact a very high false negative rate, and were wrongly attributed, in most cases, to thermals. This is reflected in the multiclass AUC, which has an average value of 0.507, a performance comparable to random chance.

**Figure S7:**
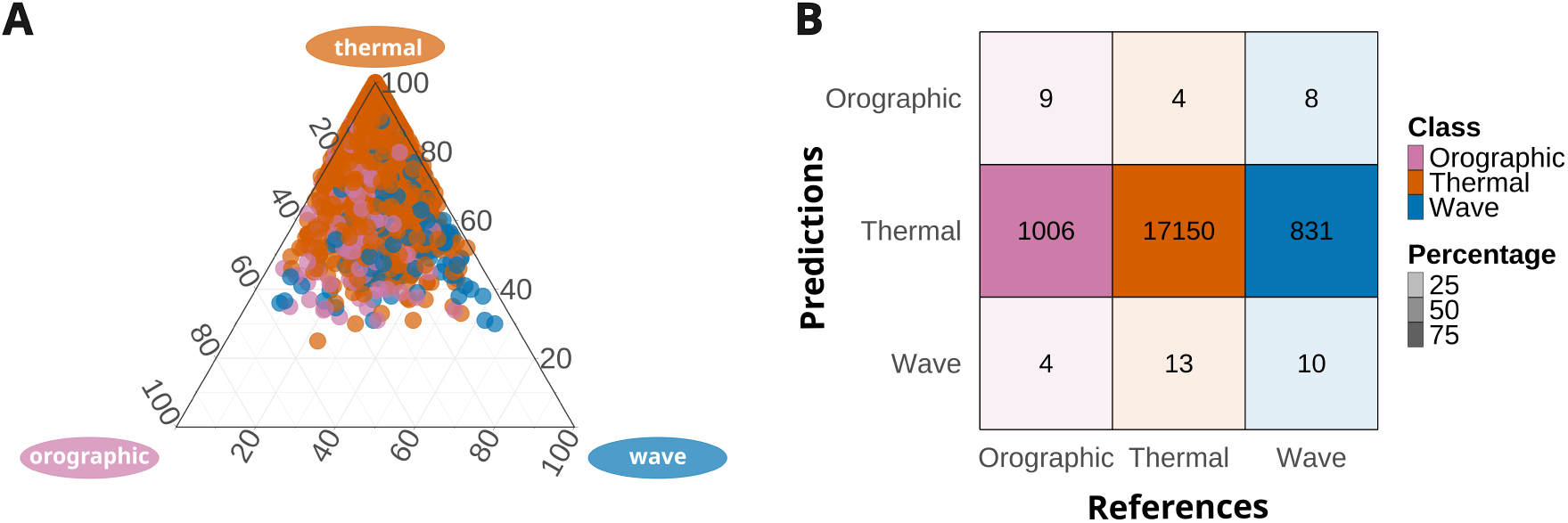
Output of the random forest models (pooled predictions from ten iterations) using behavioural predictors and frequency-weighted training dataset. The ternary plot (panel A) and the confusion matrix (panel B) show the correct classification of 9/1019 orographic uplifts, 17150/17167 thermal uplifts, and 10/849 gravity waves. The overall accuracy of the iterations is 0.902, but the model is incapable of effectively distinguish dynamic uplifts from thermal ones (multi-class AUC is 0.507).

### S7: Supplementary Tables

**Table 1:** Computed constant values for vertical wind field estimation.

| Variable | Value |
| --- | --- |
| Body Mass (mass) [kg] | 2.387-6.665 |
| Wing Span (b) [m] | 1.60-2.22 |
| Wing Area (S) [m<sup>2</sup>] | 0.5237-0.6070 |
| Air density ($\rho$) [kg m<sup>-3</sup>] | 0.957 |
| Gravitational Acceleration (g) [m s<sup>-2</sup>] | 9.81 |
| Drag Coefficient (k) | 1.1 |
| Drag Coefficient (CDo) | 0.1 |
| Bank Angle ($\Phi$) [degrees] | 25 |

**Table 2:**
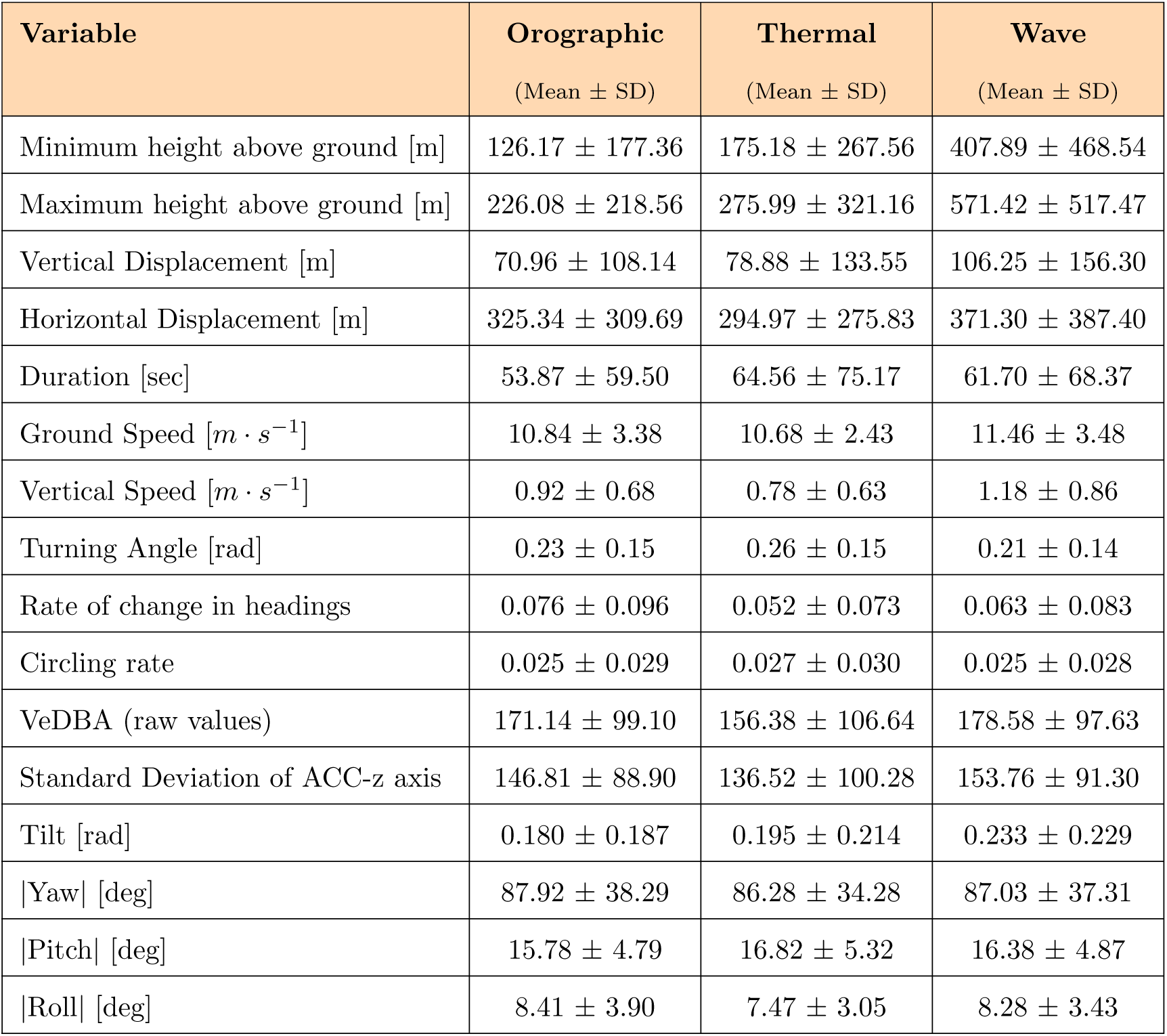
Summary of soaring flight variables by uplift type, averaged along soaring segments.

## Notes

### Competing Interest Statement

The authors have declared no competing interest.

